# EXPRESSION OF *Mycobacterium tuberculosis* RpsA IN *Mycobacterium smegmatis* INCREASES SUSCEPTIBILITY TO PYRAZINAMIDE

**DOI:** 10.1101/2020.03.31.018390

**Authors:** Ricardo Antiparra, Marco Santos, Angie Toledo, Robert H. Gilman, Daniela E. Kirwan, Patricia Sheen, Mirko Zimic

**Author notes:** Corresponding Author: Dr. Mirko Zimic, Laboratorio de Bioinformática, Biología Molecular y Desarrollos Tecnológicos. Laboratorios de Investigación y Desarrollo. Facultad de Ciencias y Filosofía. Universidad Peruana Cayetano Heredia., Av. Honorio Delgado 430, San Martín de Porres, Lima 31, Peru.

## Abstract

Pyrazinamide (PZA) is one of the most important drugs used in combined antituberculous therapy. After the drug enters *Mycobacterium tuberculosis* it is hydrolyzed by pyrazinamidase (PZAse) to the bactericidal molecule pyrazinoic acid (POA). Ribosomal protein S1 (RpsA) was recently identified as a possible target of PZA based on its binding activity to POA and capacity to inhibit trans-translation. However, its role is not completely understood.

It has been proposed that *Mycobacterium smegmatis* RpsA is not capable of binding POA, unlike *M. tuberculosis* RpsA. This may be due to the different amino acid sequence in the carboxy-terminal region of the two molecules: in *M. smegmatis* RpsA it is much closer to the sites that may interact with POA than in *M. tuberculosis* RpsA. These differences could be contributing, along with the presence of highly active POA efflux, to the natural resistance to PZA in *M. smegmatis*.

To further understand the mechanisms of action of PZA and the role of RpsA in PZA susceptibility, we evaluated the effect of complementing *M. tuberculosis* RpsA expression in *M. smegmatis* using pNIT mycobacterial non-integrative expression vector and then performed a PZA susceptibility test determining the minimum inhibitory concentration (MIC) of PZA. It was expected that chimeric ribosomes comprising *M. tuberculosis* RpsA may be present and may affect PZA susceptibility.

Our results showed a reduction in PZA MIC in *M. smegmatis* complemented with overexpressed *M. tuberculosis* RpsA compared to non-overexpressed *M. smegmatis* (468 µg/mL and >7500 µg/mL respectively).

## INTRODUCTION

Pyrazinamide (PZA) is one of the most important drugs in the treatment of tuberculosis (TB), being used in both first and second line regimens. It is uniquely effective against slowly dividing sub-populations of bacteria in TB infection, and its inclusion in first-line treatment regimens makes it possible to shorten the duration of treatment from 9 to 6 months (1). PZA susceptibility has been associated with clinical outcomes and although testing for resistance is recommended, this is not universally done due to technical challenges (2).

PZA is a pro-drug that enters *Mycobacterium tuberculosis* by passive diffusion. In the cytoplasm, PZA is converted by the enzyme pyrazinamidase (PZAse) to its active form, pyrazinoic acid (POA). The POA is then expelled out of the mycobacterium by an efflux mechanism. If the pH of the extracellular environment is acidic, the POA is protonated to POAH, re-enters the cell, and the proton is released into the cytosol. This cycle repeats itself, causing an intra-cellular accumulation of POA and a reduction in the pH of the cytoplasm. This leads to a lethal alteration of membrane permeability; however, the exact mechanism by which this occurs is not yet known (3). It is important to note that other recent studies found that extracellular acid pH is not required in order for PZA/POA achieve its lethal effect (4,5,6). This confirms that the real mechanism of action of PZA is actually more complicated than previously thought, and further research is required.

Mechanisms of resistance to PZA in *M. tuberculosis* have been mostly associated with the loss of PZAse activity due to mutations in the *pncA* gene (*pncA* mutations in >80% of clinical isolates) (7,8,9). However, there are strains of *M. tuberculosis* that are resistant to PZA yet have active PZAse. The mechanism by which these strains are resistant to PZA is not fully known, although several studies suggest the involvement of other target proteins (10,11,12).

In contrast to *M. tuberculosis, Mycobacterium smegmatis* is naturally resistant to PZA. While it actively produces POA through two PZAse enzymes (13), it was shown that in *M. smegmatis* POA is pumped into the extracellular environment 900 fold faster than in *M. tuberculosis*, preventing POA from accumulating intracellularly at a minimal required critical concentration, and consequentially from exerting any lethal effect on the mycobacterium (14).

Recent studies have identified the ribosomal protein S1 (RpsA) as a probable target of POA (15,16). RpsA is a protein involved in the ribosomal trans-translation process, which has been associated with bacterial survival in states of stress, bacterial virulence, and nutrient recovery in periods of deprivation. Blocking trans-translation prevents the recovery of stalled ribosomes and increases the accumulation of toxic or harmful proteins, causing cell death under conditions of metabolic stress (17). The same study used microcalorimetry to show that in contrast, *M. smegmatis* RpsA protein is not capable of binding POA. Even though both *M. tuberculosis* and *M. smegmatis* RpsA proteins share a high percentage of amino acid identity, some key differences exist, mainly in their terminal carboxyl region that may affect their capacity to bind to POA and thus susceptibility to PZA.

In the present study, the *Rv1630* gene encoding the *M. tuberculosis* RpsA protein was cloned in the mycobacterial non-integrative vector pNIT and expressed in *M. smegmatis* mc^2^155. PZA susceptibility was then evaluated in this genetic variant of *M. smegmatis* compared to the native strain. Our hypothesis is that the genetic variant of *M. smegmatis* would have a subpopulation of chimeric ribosomes incorporating *M. tuberculosis* RpsA able to bind POA that may inhibit trans-translation, and potentially affect the susceptibility to PZA.

## METHODS

### Bacterial strains and growth conditions

*Escherichia coli* Novablue was routinely grown in Luria Bertani (LB) medium for its use in DNA cloning procedures. *Mycobacterium smegmatis* mc^2^ 155 was grown at 37°C in Middlebrook (MB) 7H9 liquid medium or on MB 7H10 agar supplemented with 0.5% (v/v) glycerol and 0.05% (v/v) Tween 80. When required, kanamycin (20 μg/mL) was added.

### Cloning of the Rv1630 (RpsA) gene from *M. tuberculosis* H37Rv in the pNIT vector

The complete *Rv1630* gene sequence (Genbank ID: NC 000962.3) was amplified by PCR, digested, and cloned in the mycobacterial expression vector pNIT. The PCR primers contained the restriction sites for NdeI and HindIII. The primers used are listed in Table 1. Both the pNIT vector and the amplification product of the *Rv1630* gene (1446 pb) were purified and digested with NdeI and HindIII and then ligated using the DNA T4 ligase for 16 hours at 16°C. *E. coli* Novablue cells were transformed by heat shock protocol using 5 µL of the ligation reaction. After a short incubation on ice, 50 µL of chemically competent *E. coli* Novablue cells and 50 ng of DNA obtained from the ligation reaction were incubated at 42°C for 45 seconds. They were then incubated at 37°C for 1 hour and transformants were selected with kanamycin. Recombinant DNA plasmid from transformed *E. coli* Novablue were extracted using the High Pure RNA Isolation Kit (Roche) following the manufacturer’s protocol, and sequenced in sequenced in an ABI PRISM 3100 Genetic Analyzer (Applied Biosystem, Foster City, CA) in both directions.

**Table I.**
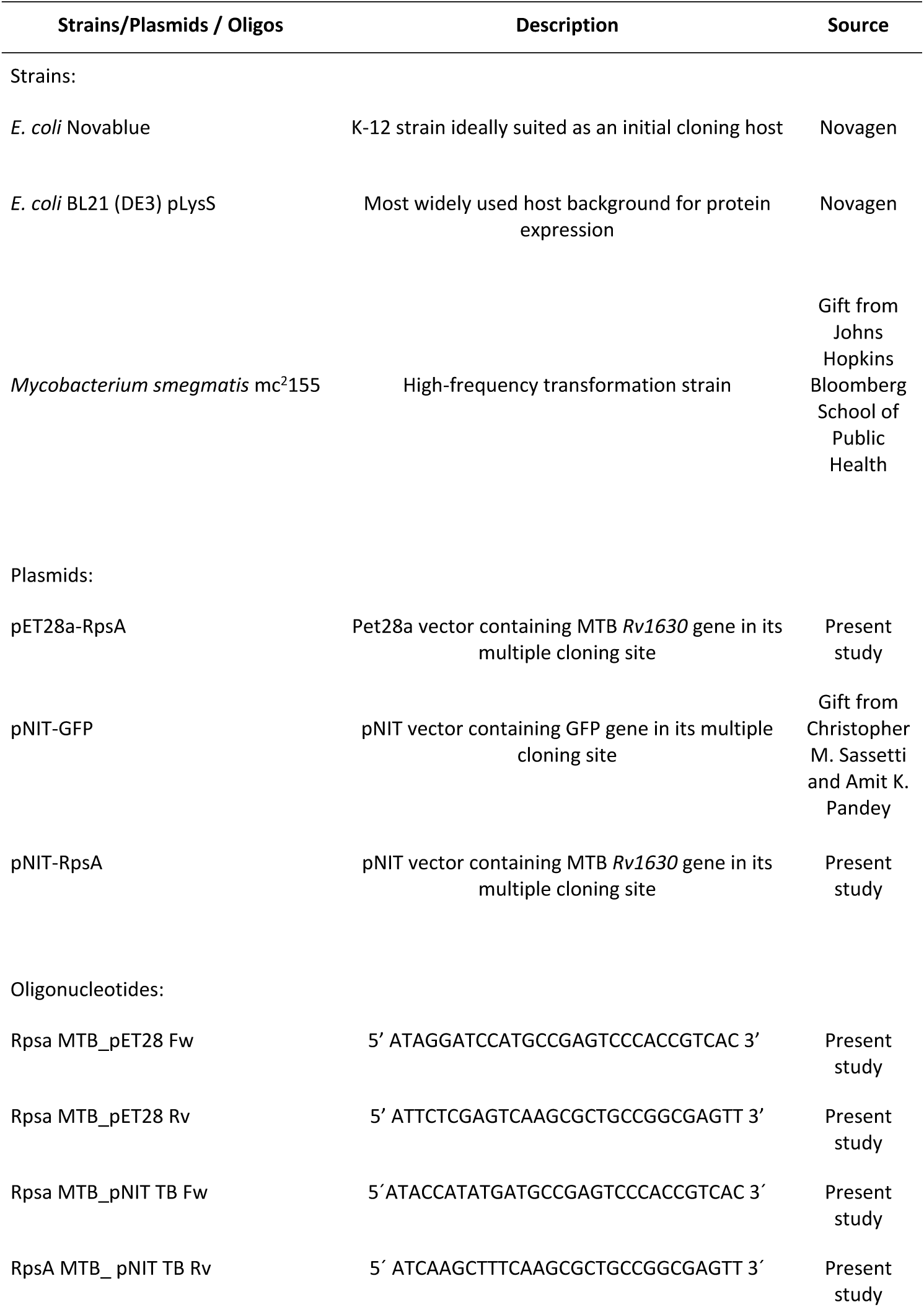
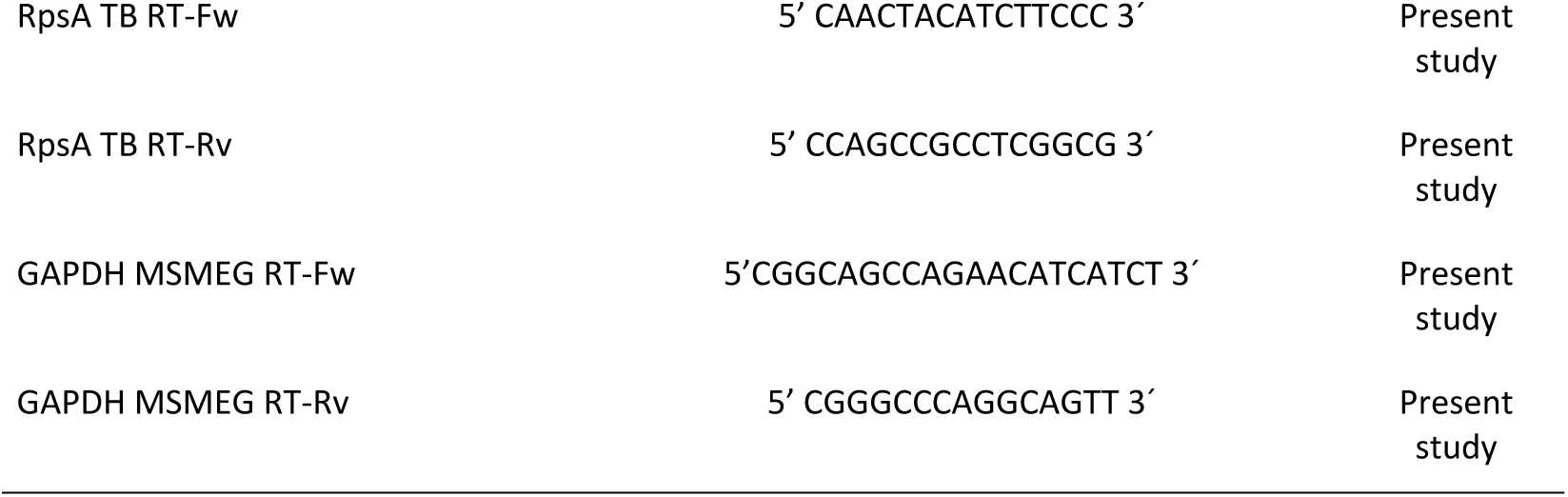
Strains, plasmids and oligos used in this study.

### Expression of the RpsA protein of *M. tuberculosis* H37Rv in *M. smegmatis* mc^2^155

*M. smegmatis* cells were grown to approximately reach an optical density measured at a wavelength of 600 nm (OD_600_) of 1, harvested, and washed 3 times in 10% glicerol-tween 80, with a reduction in the volume of 10% glycerol-tween 80 used for each wash. Finally, the washed pellet was resuspended in 1 mL of the original culture medium (20). 100 µL of electrocompetent *M. smegmatis* cells were transformed with 2 µL of the mycobacterial expression vector pNIT cloned with the gene *Rv1630* (*RpsA*) using electroporation with a gene pluser (Bio-Rad, USA). Electrocompetent *M. smegmatis* cells were pipetted into a 0.2 cm gap size cuvette with 1 µg of plasmid DNA and transformed using a setting of 2.5 kV and 1000 Ω. For the expression of the pNIT-RpsA vector, transformed *M. smegmatis* cells were induced with concentrations of 5 µM and 10 µM of Isovaleronitrile (IVN) in broth medium.

In order to verify successful expression using the pNIT vector in *M. smegmatis*, we cloned Green Fluorescent Protein (GFP) in another pNIT vector. After transforming *M. smegmatis* with pNIT-GFP we induced expression with E-caprolactam at a concentration of 10 µM. E-caprolactam was used in place of IVN because it can be incorporated into agar plates. We then detected fluorescence using UV excitation (320 nm) to confirm GFP expression.

### qRT-PCR to confirm *M. tuberculosis* Rv1630 gene transcription in *M. smegmatis*:pNIT/RpsA

*M. smegmatis* cells transformed with pNIT/ M. tuberculosis RpsA was grown at 37°C in Middlebrook 7H9 liquid medium supplemented with 0.5% (v/v) glycerol and 0.05% (v/v) Tween 80 to an OD_600_ of 0.8. After inducing with 5 µM and 10 µM of IVN, cultures were centrifuged at 12,500 rpm at 4°C for 20 minutes and the cell pellets were used for RNA extraction.

RNA extraction was performed using the Fast RNA Pro Blue kit (MP Biomedicals) on a Fast Prep-24 instrument (Applied Biosystems) following the manufacturer’s instructions. RNA quality and integrity were verified by observing the staining intensity of the major ribosomal RNA (rRNA) bands and any degradation products in a 1% agarose gel electrophoresis using TBE buffer (89 mM Tris, 89 mM boric acid, 2 mM EDTA). Absorbance at 260 nm of the extracted RNA was measured using a NanoDrop 2000 spectrophotometer (Thermo Fisher Scientific). The extracted RNA was then treated with DNAse I (RNAse free): 0.4 U of DNAse/g RNA was added to give a reaction volume of 100 µL. DNAse buffer 1X and DEPC water were then added and this was incubated at 37°C for 30 minutes followed by DNAse inactivation at 75°C for 5 min.

Reverse transcription of *M. tuberculosis Rv1630* and Glyceraldehyde-3-Phosphate Dehydrogenase (GAPDH) housekeeping gene was performed using 200 ng RNA in a T100 Thermal Cycler (Bio-Rad). The reaction was carried out with Reverse Transcription Reagents (Applied Biosystems) in a 20 µL reaction volume, which contained 4 µg of total RNA, retro-transcription buffer 1X, 5.5 mM MgCl_2_, 2 mM dNTPs, 0.5 µM of reverse primer (primers used are listed in Table 1), 0.4 U/l RNAse inhibitor, 3 U/µL Multi Scribe Reverse Transcriptase, and DEPC treated water. The following thermal parameters were used: 1 cycle to 48°C for 45 min, 1 cycle to 95°C for 5 min, and the stable phase to 5°C for 5 min. To verify that there was no genomic DNA contamination, a reaction without reverse transcriptase was added. qPCR was performed independently using specific primers that hybridize a specific sequence of the *M. tuberculosis Rv1630* (*RpsA*) gene but not the homologue gene coding for the RpsA protein of *M. smegmatis*. The primer sequences for qPCR for both the *Rv1630* and housekeeping genes are found in Table I. Values obtained were normalized with respect to the housekeeping GAPDH gene using Livak’s method (18). Each cDNA was amplified in a LightCycler Nano thermocycler (Roche) using 10 µL of LightCycler 480 SYBR Green I Master (1.6 µL of each forward and reverse primer (10 µM concentration), 2 µL of cDNA and 4.8 µL of Milliq water), under the following cycling parameters: 1 cycle of 95°C for 10 minutes, 40 cycles of 95°C for 20 seconds, 60°C for 20 seconds and 72°C for 35 seconds, 1 cycle of 60°C to 95°C with a ramp of 0.1°C/s, and 1 cycle of 40°C for 10 minutes. The cycle threshold (Ct) was calculated using Step One Plus v2.1 Software (Applied Biosystems). The experiment was repeated three times using RNA extracted from three independent cultures.

### Confirmation of RpsA protein expression by Western blot in strains of *M. smegmatis*:pNIT/RpsA

#### Cloning of M. tuberculosis H37Rv Rv1630 (RpsA) into pET28a vector

*M. tuberculosis* Rv1630 gene sequence (Genbank ID: NC 000962.3) was used to design the primers for cloning in the pET28a vector (primers used are listed in Table 1). Primers contain the restriction sites for the enzymes BamHI and XhoI and provide an amino-terminal 6His-Tag. Gene amplification was performed using a 25 µL final volume PCR master mix containing 1X PCR buffer, 1.5 mM MgCl2, 0.2 mM dNTPs, 1 µM forward and reverse primers and 0.025 U/µL of Taq polymerase. The conditions for the amplification were: an initial denaturation of 95°C for 5 min, followed by 35 cycles of a denaturation cycle at 95°C for 30 sec, an annealing cycle at 62°C for 1 min, and an extension cycle at 72°C for 90 sec. A final extension cycle was performed at 72°C for 5 min. Both the pET28a vector and the *Rv1630* gene amplification product (1446 bp) were purified using the PCR quick spin kit (Roche) following the manufacturer’s protocol. They were then digested with BamHI and XhoI and inserted into the pET28a vector using the DNA T4 ligase for 16 hours at 16°C. The *E. coli* Novablue cells were transformed by thermal shock treatment using a volume of 5 µL of the ligation reaction. Plasmid DNA was extracted from recombinant clones with the High Pure Isolation kit (Roche) as directed by the manufacturer and sequenced in an ABI PRISM 3100 genetic analyzer (Applied Biosystem, Foster City, CA) in both directions.

### Expression in *E. coli* and purification of *M. tuberculosis* H37Rv recombinant RpsA

*E. coli* cells BL21 (DE3) pLysS were transformed by thermal shock treatment using 1 µL of the expression vector pET28a cloned with the *M. tuberculosis* Rv1630 gene. Subsequently, transformed colonies were inoculated in 50 ml of Luria Bertani (LB) broth at 37°C overnight. The culture was incubated at 37 degrees for approximately 2 hours until reaching an OD_600_ of 0.6-0.8. 500 µL of Isopropyl β-D-thiogalactoside (IPTG) was then added at a final concentration of 0.5 mM and this was incubated for 4 hours at 37°C. The cells were separated by centrifugation at 6,000 rpm at 4°C for 5 min. The pellet was resuspended in 30 mL of binding buffer (20 mM Imidazol, 0.5 M NaCl and 20 mM phosphate buffer, pH 7.4) and lysed by repeated thermal shock cycles (−70°C to 37°C) followed by a sonication phase using an Ultrasonic Liquid Processor Homogenizer (Misonix S3000, 3 series of 0.1 sec ON / 0.1 sec OFF for one minute). After centrifugation at 12,500 rpm at 4°C for 20 minutes, the supernatant was purified by affinity chromatography using a 5 mL His-trap column that binds 12 mg/mL of proteins. The column was then washed with 25 ml binding buffer. RpsA protein bound to the column was eluted using pH 7.4 phosphate buffer with 100 mM imidazole. The 5 ml fractions obtained were combined and concentrated 20 times by filtration with PBS (137 mM NaCl; 2.7 mM KCl; 4.3 mM Na2HPO4; 1.47 mM KH2PO4), for which a cellulose membrane (10 kDa) was used in the Amicon Ultracel (Millipore) ultrafiltration system at 4°C. The protein concentration was determined by Bradford’s reaction, in which a standard curve was constructed using Bovine Albumin Serum (BSA).

### Ethics statement, animals and immunization protocol for the production of rabbit anti-RpsA hyperimmune polyclonal antibodies

Animal experiments were carried out with strict adherence to the Ethics Commission on Animal Use (CEUA) from the Peruvian University Cayetano Heredia. Two 4-month-old rabbits were immunized four times with 1 mg of recombinant *M. tuberculosis* RpsA protein using Freund’s complete adjuvant (FCA). Pre-immune and post-immune serum samples were obtained by collecting approximately 3 mL of venous blood obtained from the rabbits’ ears. After the fourth immunization, 75 days post initial immunization, a final bleed was performed by cardiac puncture under anesthesia (60 mg/kg Ketamine, Ket-A-10, Alfasan; and 40 mg/kg Xylazine, Xyl-A-2, Alfasan) to collect an approximate volume of 30 mL of blood. Once anesthetized, the rabbits were sacrificed with an overdose of Halatal at 20 mg/kg which was administered intramuscularly.

An ELISA assay was performed to evaluate whether the post-immune serum contained polyclonal anti-RpsA antibodies. 100 µL of recombinant RpsA (1 µg/mL, 5 µg/mL, and 10 µg/mL) was fixed in bicarbonate-carbonate buffer at 4°C overnight on an ELISA microplate (INMULON Plates). The microplates were washed with 0.05% PBS-Tween 20 (PBS-T), and each specific coated microplate was blocked using 3% bovine serum albumin (BSA) and incubated at 37°C for 1 hour. The microplates were then washed 5 times with PBS-T, 0.05%, 200 µL per well. Subsequently, the post immune serum was diluted with 0.05% PBS-T and 100 µL was added to each of the wells and incubated at 37°C for 1 hour in a wet chamber environment. The microplates were then washed 5 times with 200 µL PBS-T, 0.05%. Subsequently, 50 µL of a 1:50,000 dilution of HRP anti-rabbit conjugate IgG in 5% skimmed milk powder was added to each well and incubated at 37°C for 1 hour. The microplates were washed 5 times with 200 µL PBS-Tween 20, 0.05%. Finally, 50 µL of OPD in citrate-phosphate buffer with sodium perborate was added to each well and incubated for 15 minutes. 50 µL of 1.25 M of H_2_SO_4_ was added to each well to stop the reactions, and absorption was measured at 595 nm in an ELISA reader (Spectramax, Molecular Devices).

### Western Blotting

Western blot against the *M. tuberculosis* RpsA protein was performed to confirm expression of RpsA protein in strains of *M. smegmatis*: pNIT/RpsA. *M. smegmatis* cultures with two concentrations of IVN inducer (5 and 10 µM) were resolved in a 12% SDS-PAGE and the proteins transferred to nitrocellulose membranes. Membranes were washed with 0.3% PBS-Tween and incubated at 4°C for 16 hours with the primary hyperimmune polyclonal anti-RpsA antibody diluted in 3% BSA. Membranes were washed 6 times with 0.3% BSA-Tween and incubated at 37°C for 1 hour with the secondary rabbit anti-IgG antibody marked with Horseradish Peroxidase (HRP). Membranes were washed and revealed with hydrogen peroxide. Finally, the formation of the immunocomplexes was visualized using the substrate diaminobenzidine (1 mg/mL in citrate buffer, pH 5.0, containing 0.05% H_2_O_2_).

### PZA susceptibility testing of induced and uninduced *M. smegmatis*:pNIT/RpsA strains using Tetrazolium Microplate Assay (TEMA)

Susceptibility to PZA was determined by minimum inhibitory concentration (MIC) measurement using the Tetrazolium Microplate Assay (TEMA) (19) for both the induced (10 µM of IVN) and uninduced *M. smegmatis*: pNIT/RpsA cultures. Final well PZA concentrations used were 7500 µg/mL, 3250 µg/mL, 1870 µg/mL, 937 µg/mL, 468 µg/mL, 234 µg/mL, and 117 µg/mL. A positive control well containing bacterial suspension without PZA was included. 7H9 broth was enriched with OADC (Oleic acid, albumin, dextrose and catalase) at pH 6. The plates were incubated for 2 days at 37°C. After growth in the positive control well on the second day of incubation, 50 µL of MTT tetrazolium salt (1 mg/mL) was added to the well and the plate was incubated for 3 hours. If a yellow to violet color change occurred, this was taken to indicate the viability of *M. smegmatis* and therefore the remaining wells were revealed at this time. MIC was defined as the lowest concentration of the drug where no color change occurred.

## RESULTS

### Cloning of *M. tuberculosis* H37Rv Rv1630 into the pNIT vector

PCR of transforming colonies following the ligation process between the pNIT vector and the *M. tuberculosis* Rv1630 gene confirmed the presence of the gene (Figure 1).

**Figure 1.**
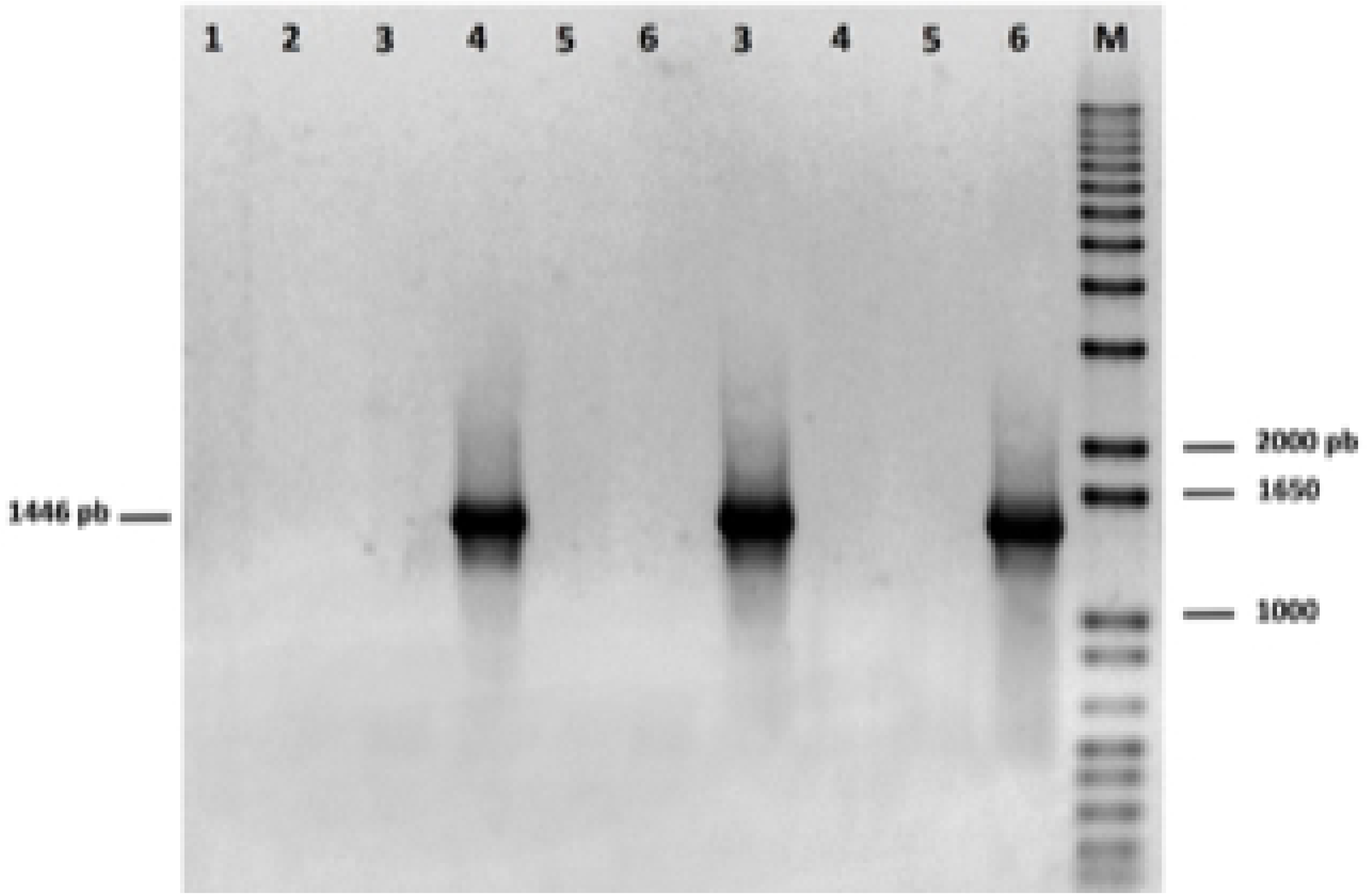
PCR Colony used to verify the insertion of the *Rv1630* gene (*M. tuberculosis RpsA*) in the pNIT vector. *Rv1630* gene: 1446 pb. Line 1-10: C1-C10. M: 1Kb plus marker (invitrogene). C: Clone.

### Expression of Green Fluorescent Protein into the pNIT vector

Visualization of GFP fluorescence in the *M. smegmatis* pNIT-GFP confirmed that the expression vector is producing the recombinant protein appropriately (Figure S1 supplementary material).

### qRT-PCR confirmation of the expression of *M. tuberculosis* Rv1630 in uninduced and induced strains of *M. smegmatis*:pNIT/RpsA

The uninduced *M. smegmatis* pNIT/ *M. tuberculosis* RpsA strain had a Ct value of 27.36 for the *Rv1630/RpsA M. tuberculosis* gene, while Ct was 13.52 for the induced culture. Ct values for the housekeeping GAPDH gene were 24.63 and 22.24 for the *M. smegmatis* pNIT/*M. tuberculosis* RpsA strain without induction and for the induced strain, respectively (Table II). The normalized expression of the *Rv1630* gene with respect to the housekeeping GAPDH gene resulted in a higher level of expression in the induced strain (Ct-RpsA/Ct-GAPDH = 13.52/22.4 = 0.60) compared to the uninduced strain (Ct-RpsA/Ct-GAPDH = 27.36/24.63 = 1.11) (P=0.046, Wilcoxon Rank Test).

**Tabla II.** Expression of *Rv1630* gene in *Mycobacterium smegmatis* mc^2^155 as determined by qPCR Real-time.

| Gene | Cycle threshold (Ct) values |  |  |
| --- | --- | --- | --- |
| | Mean CT<br>(MSMEG # Rv1630 IND)<sup>a</sup> | Mean CT<br>(MSMEG # Rv1630 NON-IND)<sup>b</sup> | ΔCT<br>(Rv1630 IND &<br>Rv1630 NON-IND) |
|  | (A) | (B) | (B)-(A) |
| *Rv1630* (*M. tuberculosis* RpsA) | 13.52 | 27.36 | 13.84 |
| GAPDH | 22.24 | 24.63 | 2.39 |
| Relative nomalized difference in expression ($\Delta\Delta$CT) | | | 11.45 |

<sup>a</sup> *M. smegmatis* mc<sup>2</sup>155 containing induced recombinant vector (MSMEG-pNIT-Rv1630); <sup>b</sup> *M. smegmatis* mc<sup>2</sup>155 containing non-induced recombinant vector (MSMEG-pNIT-Rv1630); GAPDH was used as reference gene; MTB, *Mycobacterium tuberculosis*.

### Confirmation of RpsA protein expression by Western blot in strains of *M. smegmatis*:pNIT/RpsA

Recombinant clones were confirmed by the release of fragment Rv1630 from vector pET28a after digestion with appropriate restriction enzymes (Figure 2). Seven recombinant clones were obtained (Vector pET28a/Rv1630). The induced fraction confirmed the presence of the RpsA protein of *M. tuberculosis* with an estimated weight of 61.2 KDa using SDS-PAGE (Figure 3).

**Figure 2.**
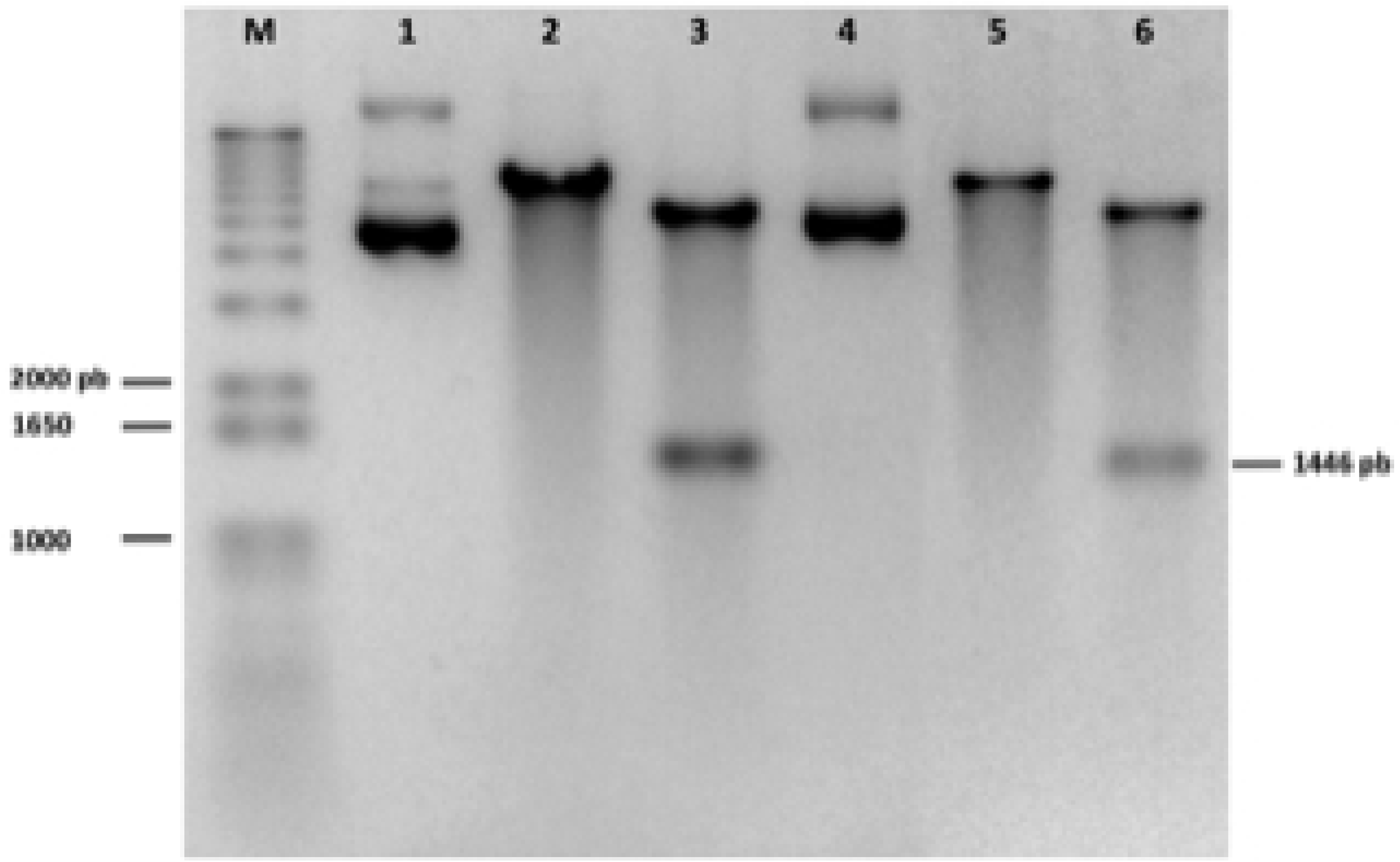
Electrophoresis in 1 % agarose gel stained with ethidium bromide. Enzymatic digestion used to test the insertion of the *Rv1630* gene (*M. tuberculosis RpsA*) in the pET28a vector. *Rv1630* gene: 1446 pb. M: 1Kb plus marker (invitrogene). Line 1: Non-digested Plasmid (Clone 1), Line 2: Plasmid + BamHI, Line 3: Plasmid + BamHI/XhoI, Line 4: Non-digested Plasmid (Clone 2), Line 5: Plasmid + BamHI, Line 6: Plasmid + BamHI/XhoI.

**Figure 3.**
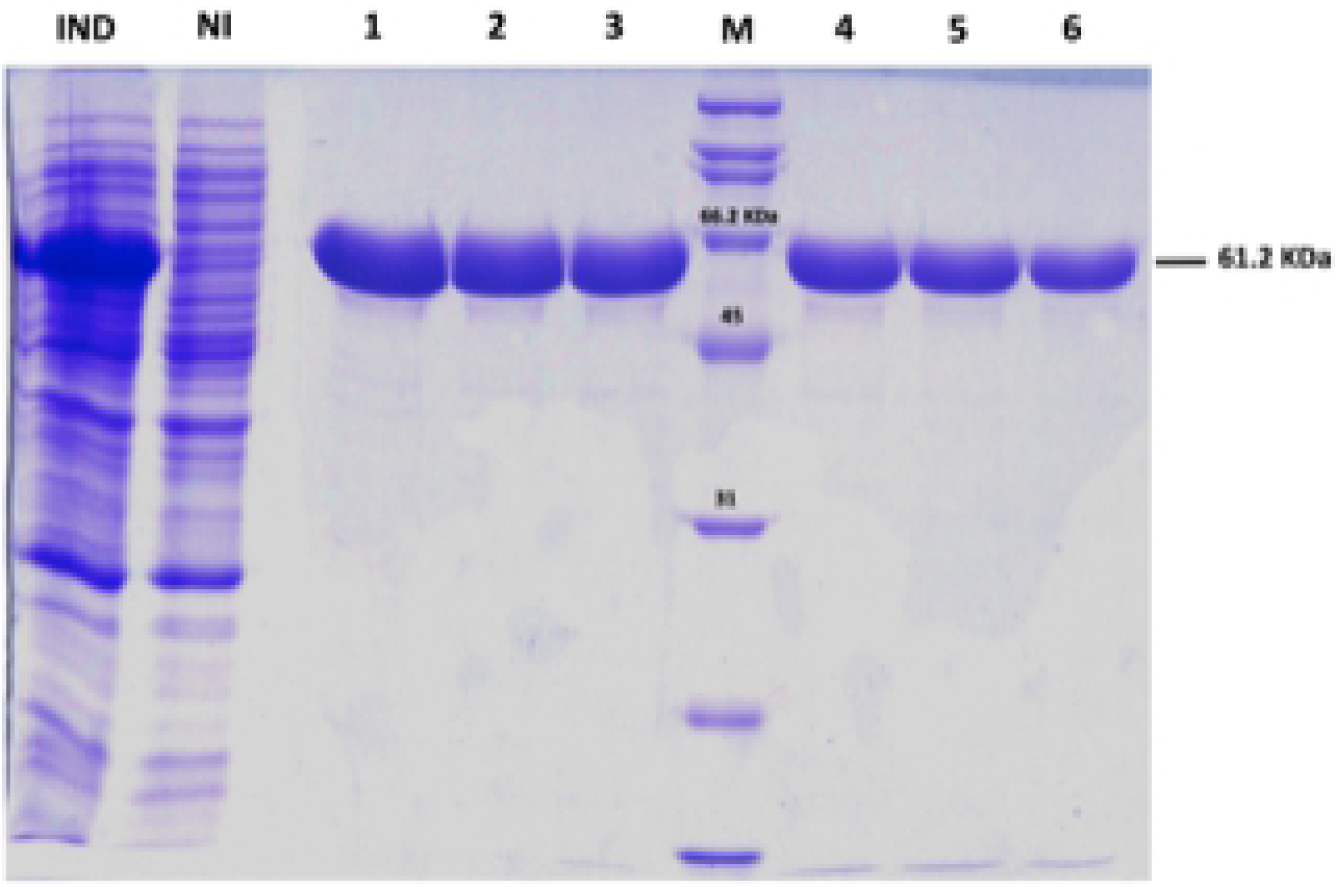
Electrophoresis SDS-PAGE 12% (SDS-PAGE). Expression and purification of the RpsA protein (*M. tuberculosis* Rv1630) in *E. coli* using the pET28a expression vector. IND: Crude extract (induced sample), NI: Crude extract (Non-induced sample), Line 1-3: Purified fractions (100 mM Imidazol), M: Broad Range protein marker (Biorad), Line 4-6: Purified fractions (100 mM Imidazol).

IVN inductor at a concentration of 10 µM enabled overexpression of *M. tuberculosis* RpsA in *M. smegmatis* pNIT/RpsA. RpsA was detected on Western blot as a 61.2-kDa band in the induced culture (Figure 4) and therefore RpsA is predicted to be a ∼61.2-kDa protein.

**Figure 4.**
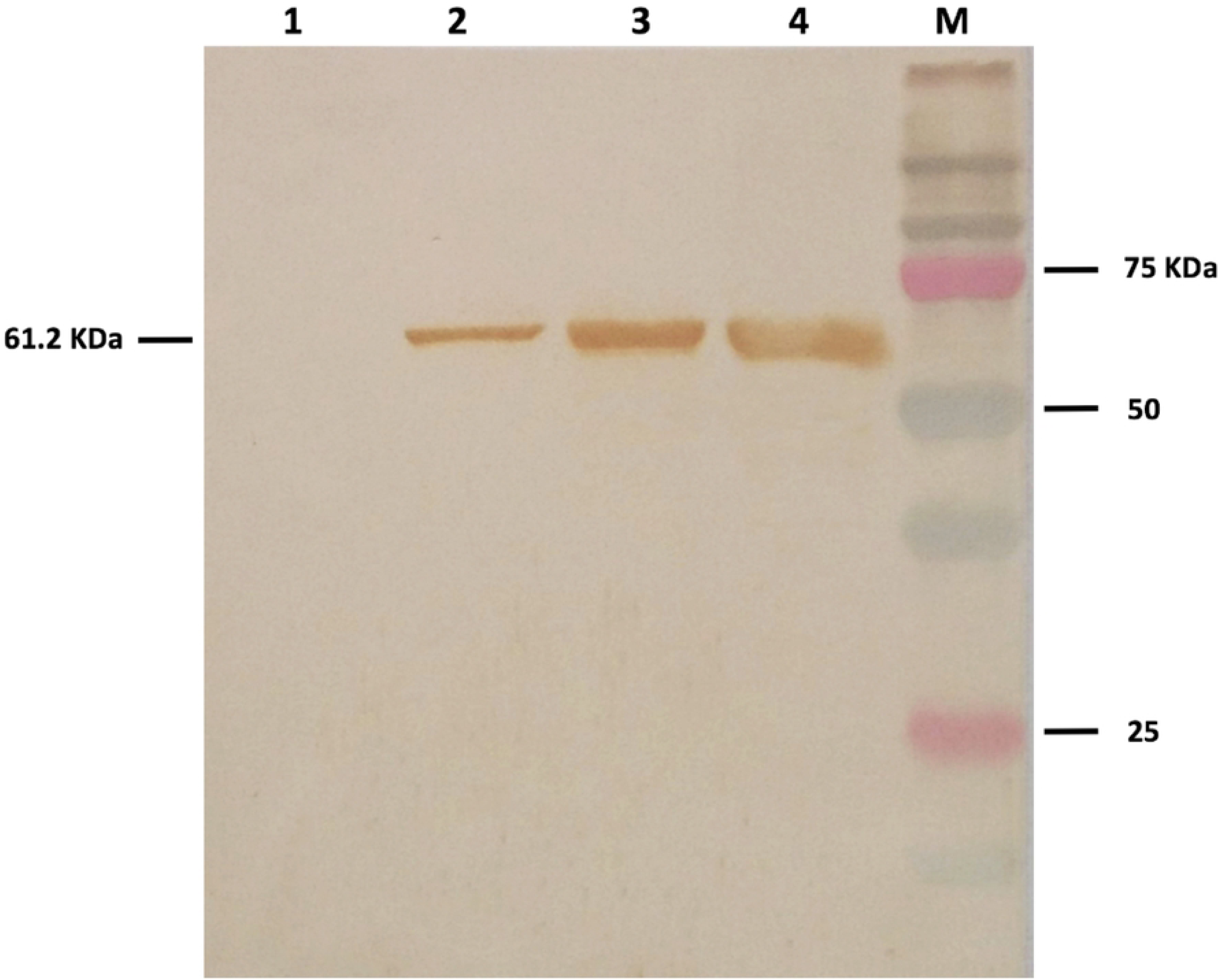
Western Blot against the *M. tuberculosis* protein RpsA. Line 1: Empty; Line 2: Culture of Non-induced *M. smegmatis* mc^2^155. Line 3: Culture of *M. smegmatis* mc^2^155 induced with 5 µM of IVN. Line 4: Culture of *M. smegmatis* mc^2^155 induced with 10 µM of IVN. M: Dual Color protein marker (Biorad). Estimated size of the *M. tuberculosis* protein RpsA: 61.2 Kda.

### PZA susceptibility testing of induced and uninduced *M. smegmatis*:pNIT/RpsA strains using TEMA (Tetrazolium Microplate Assay)

The TEMA assay confirmed a growth difference between the culture of uninduced versus induced *M. smegmatis* pNIT/RpsA in the presence of PZA, whereby there is a notable inhibition in the growth of induced *M. smegmatis* pNIT/RpsA cultures which express *M. tuberculosis* RpsA (Figure 5).

**Figure 5.**
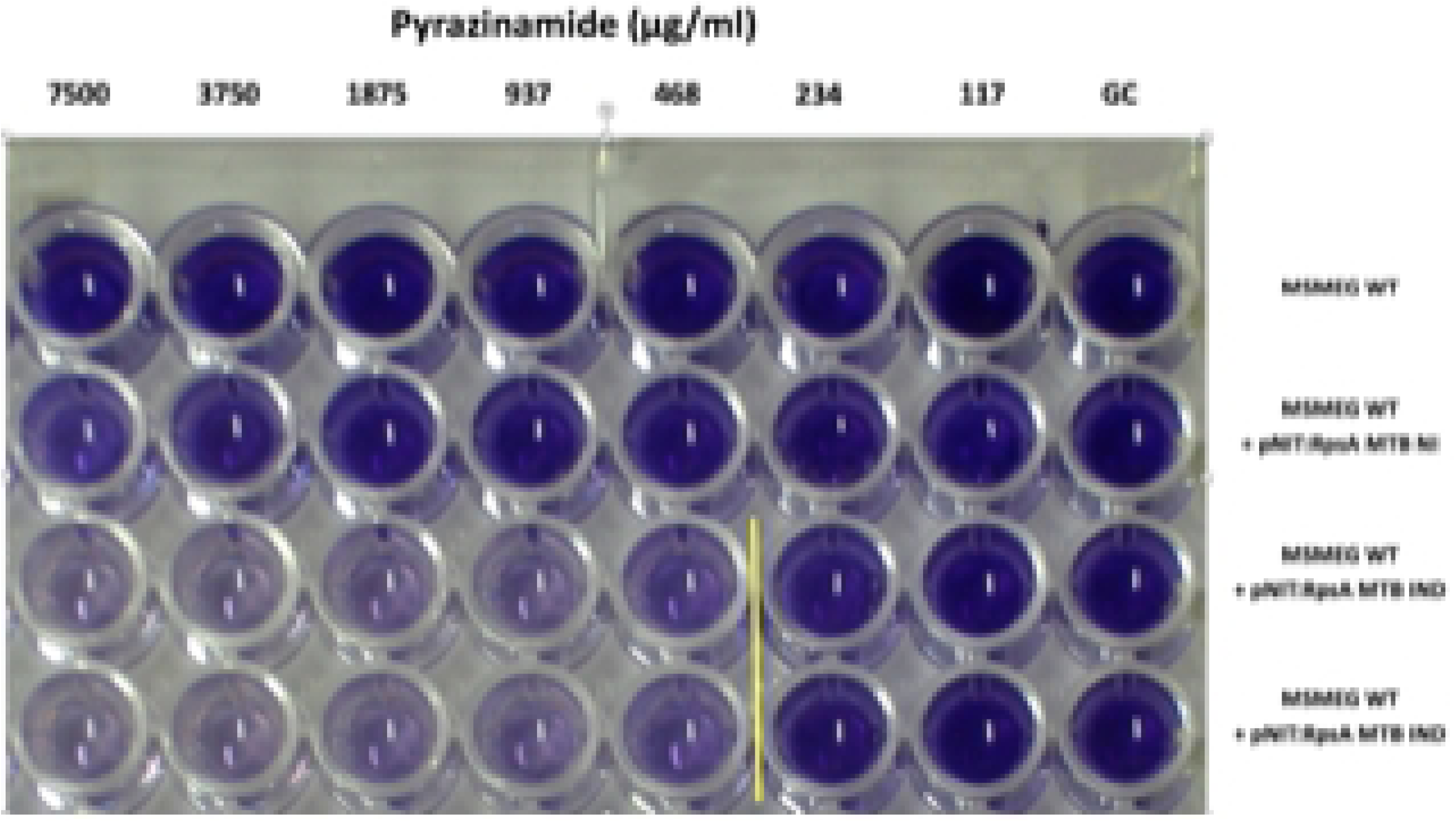
MIC of PZA in cultures of *M. smegmatis* using the TEMA assay. Line 1, *M. smegmatis* mc^2^155 WT without plasmid. Line 2, *M. smegmatis* mc^2^155 pNIT:RpsA Non-induced plasmid. File 3, *M. smegmatis* mc^2^155 with pNIT:RpsA plasmid induced with 10 µM IVN. Row 4, *M. smegmatis* mc^2^155 with pNIT:RpsA plasmid induced with 10 µM IVN (duplicate). PZA serial dilutions from 7500 µg/ml to 117 µg/ml are shown. GC: Positive growth control in a drug-free 7H9 Broth.

Evaluations in triplicate on different days show an average MIC reading that confirms a change from a naturally PZA resistant phenotype in *M. smegmatis* (MIC: >7.5 mg/mL) to a phenotype susceptible to PZA (MIC: 0.468 mg/mL) in the induced *M. smegmatis* mc^2^155 pNIT:RpsA *M. tuberculosis* RpsA (P=0.025, Wilcoxon Rank Test) (Table III).

**Table III.** MIC of pyrazinamide in *M. smegmatis* mc^2^155 strains harbouring foreign DNA *(M. tuberculosis Rv1630*)

| Drug | MIC (µg/ml) |  |  |
| --- | --- | --- | --- |
|  | *M. smegmatis* mc<sup>2</sup>155 | *M. smegmatis* mc<sup>2</sup>155 (pNIT-Rv1630 NON-IND)<sup>a</sup> | *M. smegmatis* mc<sup>2</sup>155 (pNIT-Rv1630 IND)<sup>b</sup> |
| PZA | > 7500 | > 7500 | 468 |

<sup>a</sup> *M. smegmatis* mc<sup>2</sup>155 containing induced recombinant vector (MSMEG-pNIT-MTB Rv1630);
<sup>b</sup> *M. smegmatis* mc<sup>2</sup>155 containing non-induced recombinant vector (MSMEG-pNIT-MTB Rv1630); PZA, Pyrazinamide; MIC, minimum inhibitory concentration.

## DISCUSSION

In contrast to *M. tuberculosis, M. smegmatis* is naturally resistant to PZA and is therefore a useful model in which to study PZA resistance mechanisms (20). This study evaluated the effect of introducing *M. tuberculosis* RpsA into *M. smegmatis* on PZA sensitivity. We observed a decrease in MIC from >7500 µg/mL in the uninduced *M. smegmatis:*pNIT:RpsA strain to 468 µg/mL when this strain was induced to produce *M. tuberculosis* RpsA, resulting in a more sensitive phenotype.

*M. tuberculosis* RpsA was expressed in *M. smegmatis* using the expression vector pNIT (induced by IVN 10 µM) which shows inducible regulation capacity and a low level of basal expression (21). Western Blot analysis verified that a greater quantity of *M. tuberculosis* RpsA was observed in the induced than in the uninduced *M. smegmatis* strain. This is similar to previous findings following expression of the *M. tuberculosis* protein PknA in *M. smegmatis* (22). The pNIT vector has also previously been used to perform functional complementation tests; for example, Li *et al*. were able to evaluate the importance of the mutant protein PknA K42M for growth in *M. smegmatis* (23). Another physiological study used pNIT to evaluate the role of PstP phosphatase proteins in the growth of the pathogen within the host. In this study, overexpression of the *PstP* gene in *M. smegmatis* led to an elongation of the cells (24).

Given that the RpsA proteins of *M. tuberculosis* and *M. smegmatis* share over 90% of their identity, it is likely that when *M. smegmatis* expresses the foreign RpsA protein from *M. tuberculosis*, a heterologous complementation process within the ribosome takes place generating groups of both native and hybrid/chimeric ribosomes. There have been studies in which the feasibility of this process has been evidenced; for example, heterologous complementation of yeast ribosomes with ribosomal proteins from plants and mammals (25,26), of fungi with human ribosomal proteins (27), and even of ribosomes from organisms of different domains such as *E. coli* and rats (28,29) have been analyzed using this same approach. In these latter cases it is important to highlight that hybrid ribosomes acquire certain functional characteristics of eukaryotic ribosomes, and therefore it is expected in our study that hybrid ribosomes in *M. smegmatis* would acquire characteristics of a ribosome from *M. tuberculosis*. In addition, there are studies in which heterologous complementation of other enzymes has been analysed (30, 31, 32, 33); for example, Konse-Thomas *et al*. evaluated the RNA polymerase synthesis activity of different RNA polymerase hybrids formed from subunits from *E. coli* and *S. marcescens* (30).

Our previous studies showed that there is an association between PZA resistance and POA efflux rate, which correlates with extracellular POA concentration (11). In *M. smegmatis*, resistance to PZA is associated to a POA efflux rate 900 times faster than that of *M. tuberculosis*, which prevents its intracellular accumulation and acidification of the cytosol and thus contributing to resistance to PZA. In a study performed by Boshoff *et al*. the gene encoding one of the two *M. smegmatis* pyrazinamidases (*pzaA*) was overexpressed and this led to a decrease in the MIC from >2,200 µg/mL to 150 µg/mL (34). This suggests that the production of POA at high levels due to PZAse overexpression leads to sufficient accumulation in the cytoplasm to cause a change in PZA susceptibility of the organism. Similarly, our results suggest that accumulation of free *M. tuberculosis* RpsA would enable the retention of intracellular POA, affecting PZA susceptibility. Consequently, the difference in amino acid composition of *M. smegmatis* RpsA compared to the recombinant *M. tuberculosis* RpsA may contribute to the natural resistance of *M. smegmatis* to PZA. This inference is consistent with the proposed mechanism for PZA resistance in *M. tuberculosis* described by Shi *et al*. (2019) (35). The trans-translation mechanism, which involves the RpsA protein, is more important in slow-growing mycobacteria (i.e. *M. tuberculosis*) than in fast-growing mycobacteria (i.e. *M. smegmatis*) where it is considered to be dispensable. However our results showed that expression of the *M. tuberculosis* ribosomal protein in *M. smegmatis* led to inhibition of growth in the presence of PZA, suggesting that trans-translation may indeed also be of importance in these mycobacteria. Further studies are needed to evaluate recombinant *M. tuberculosis* RpsA in hybrid/chimeric ribosomes during trans-translation in *M. smegmatis* and its effect on PZA MIC reduction. These data may then be used to infer the relationship between RpsA structure and function and PZA resistance in *M. tuberculosis*.

## ETHICAL STATEMENT

Ethical approval for rabbit serum collection was obtained from the institutional Ethics Committee of Universidad Peruana Cayetano Heredia with code 66397 (CIEA-Session 09 March 2016).

## ACKNOWLEDGEMENTS

We thank Christopher M. Sassetti and Amit K. Pandey for their kind gifts of pNIT-GFP.

## FIGURE LEGENDS

**Figure S1.**
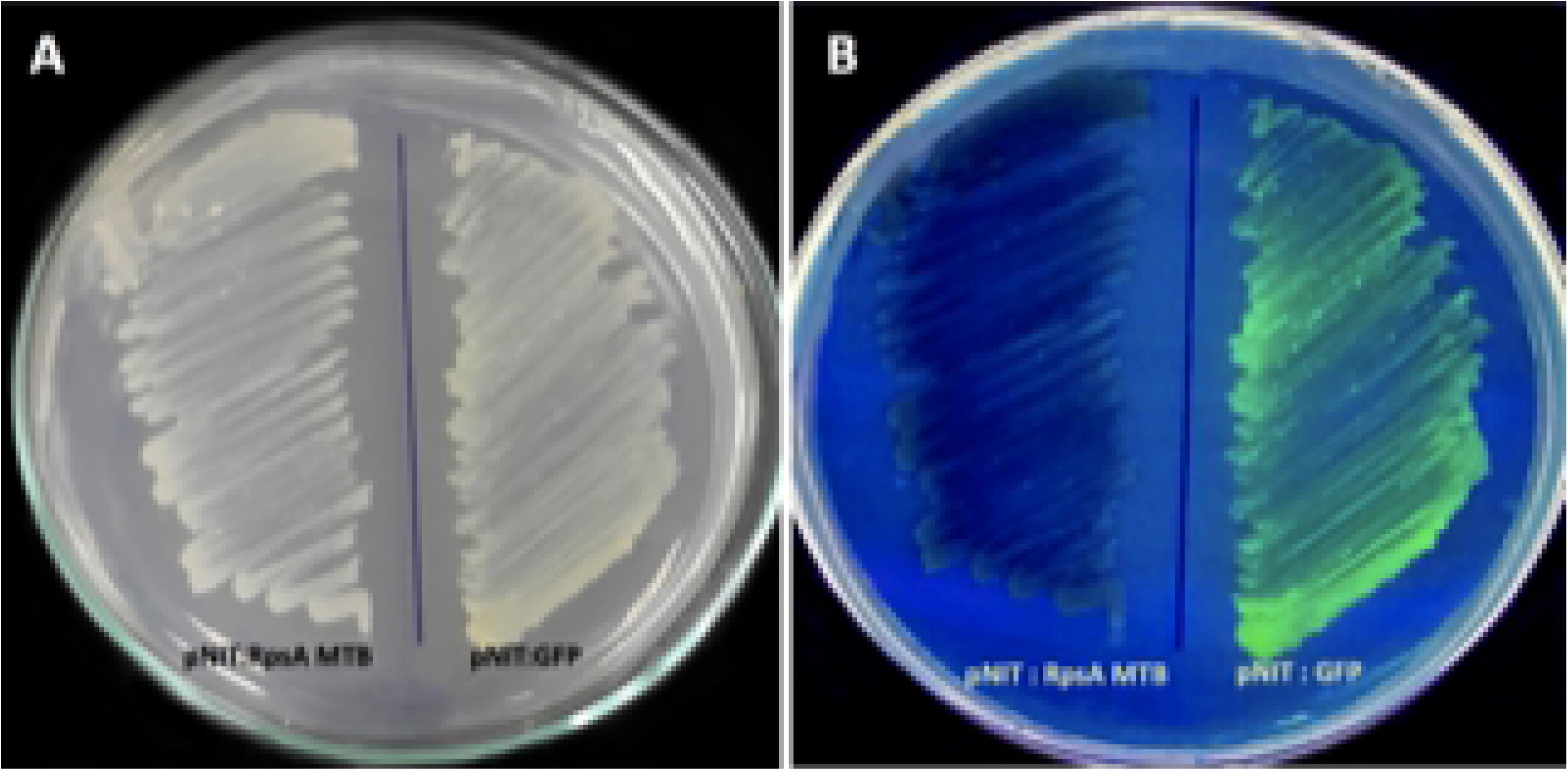
SUPPLEMENTARY MATERIAL. Cultures of *M. smegmatis* mc^2^155 with recombinant vectors pNIT/RpsA and vector pNIT/GFP without exposure to UV light (A) and with exposure to UV light (B).

